# Spatial transcriptomic profiling of human paravertebral sympathetic chain ganglia reveals diabetes-induced neuroplasticity

**DOI:** 10.1101/2025.06.24.661373

**Authors:** Marisol Mancilla Moreno, Pooja J. Patel, Kree Goss, Subhaan M. Mian, Sera Nakisli, Nikhil Inturi, Stephanie I. Shiers, Peter Horton, Anna Cervantes, Geoffrey Funk, Tariq Khan, Erin Vines, Rebecca M Klein, Darrell A Henze, Diana Tavares-Ferreira, Theodore J Price, Muhammad Saad Yousuf

**Affiliations:** Center for Advanced Pain Studies and Department of Neuroscience, School of Behavioral and Brain Sciences, University of Texas at Dallas, Richardson, TX 75080; Southwest Transplant Alliance, Dallas, TX 75231; Department of Neuroscience, Merck & Co., Inc., Rahway, NJ 07065 USA

**Keywords:** sympathetic chain ganglia, spatial RNA sequencing, diabetes, dorsal root ganglia, drug development

## Abstract

The paravertebral sympathetic chain ganglia (SCG) are autonomic ganglia critical for regulating the “fight-or-flight” response. Symptoms of sympathetic dysfunction are prevalent in diabetes, affecting up to 90% of patients. The molecular and cellular composition of the human SCG and its alteration in diabetes remains poorly defined. To address this gap, we performed spatial transcriptomic profiling of lumbar SCGs from diabetic and non-diabetic organ donors. We identified 3 three distinct neuronal populations, two noradrenergic (NA1 and NA2) and one cholinergic (CHO), based on tyrosine hydroxylase (*TH*) and *SLC18A3* expression, respectively. We also characterized 9 non-neuronal populations consisting of Schwann cells, immune cells, fibroblasts, adipocytes, and endothelial cells. In diabetic SCGs, we observed a significant loss of myelinating Schwann cells and a phenotypic shift of cholinergic neurons toward a noradrenergic identity. Additionally, diabetes was associated with a significant reduction in the transcripts of vasodilatory neuropeptides, such as *VIP* and *CALCA*, suggesting a mechanism for impaired vascular control. Upstream regulator analysis highlighted altered neurotrophic signaling in diabetes, with enhanced NGF/TRKA and diminished BDNF/TRKB activity, potentially driven by target-derived cues. Comparison between SCG and dorsal root ganglia (DRG) neurons revealed ganglia-specific genes, like *SCN3A* and *NPY* (SCG) versus *SCN10A* and *GPX1* (DRG), offering specific therapeutic targets for autonomic dysfunction or pain. Our findings provide a transcriptomic characterization of human SCG, revealing molecular signatures that underlie diabetic autonomic dysfunction. This work lays a foundation for the development of therapies to restore sympathetic function and avoid unintended autonomic effects in the development of analgesics.

**Significance Statement:** Autonomic dysfunction affects up to 90% of people with diabetes, yet the human sympathetic nervous system remains poorly molecularly defined. To address this gap, we present a spatial transcriptomic profile of the human sympathetic chain ganglia (SCG), revealing how diabetes affects the human autonomic nervous system. We show that diabetes shifts the cholinergic neuronal population to a noradrenergic phenotype and reduces vasodilation neuropeptide expression, potentially explaining impaired vascular control and thermoregulation. Comparative analysis of sympathetic and sensory ganglia reveals distinct gene profiles that may inform novel therapeutic strategies. These findings offer critical insight into the molecular drivers of diabetic autonomic neuropathy and lay the groundwork for safer, more precise treatments that selectively modulate autonomic or sensory function in chronic disease.

## Introduction

Diabetes mellitus is a chronic metabolic disorder that is characterized by persistent hyperglycemia. It is associated with multiple system-wide complications, like autonomic dysfunction, that significantly impair patients’ quality of life and increase healthcare burden. Autonomic dysfunction affects up to 90% of diabetic patients and consists of cardiovascular, gastrointestinal, genitourinary, metabolic, and sudomotor symptoms (1, 2). The autonomic nervous system (ANS) consists of three divisions, the sympathetic (SNS), the parasympathetic, and the enteric nervous systems, which collectively regulate involuntary bodily functions (3). Disruption in sympathetic signaling due to diabetes has been increasingly recognized as a contributing factor in the development and progression of diabetic complications, like heart disease, hypertension, nephropathy, and sensory neuropathy (2).

Anatomically, the paravertebral sympathetic chain ganglia (SCGs) are arranged bilaterally along the spinal column, forming a chain of interconnected ganglia that extends from the base of the skull to the coccyx (4). Each ganglion receives preganglionic input from the spinal cord and sends postganglionic axons to all organ systems (5). Preganglionic neurons are located in the intermediolateral nucleus of lamina VII of the T1-L2 spinal cord segments which go on to synapse onto postganglionic neurons of the SCG on either the same level or a superior or inferior ganglion of the chain (6). Postganglionic neurons of the SCG are terminal mediators of the cardiovascular tone of target organs and thermoregulation (6). The SCG’s unique organization allows for coordinated and efficient transmission of sympathetic signals, ensuring rapid responses to physiological demands.

Despite the widespread and profound effect of diabetes on the SNS, the molecular architecture and cellular complexity of the human SCG is not well defined. Prior reports, as early as 1885, characterizing the anatomy of human SCG have been biased by the identification of select neuronal markers of neurotransmitters, such as norepinephrine (NE), and neuropeptides, like neuropeptide Y (NPY) (7). Recent efforts have been concentrated towards delineating cellular identities in rodent SCGs. Using single cell transcriptomics, Furlan, *et al*. (8) and Mapps, *et al*. (9) identified unique neuronal and non-neuronal populations in mouse SCG which were distinct from their sensory counterparts in the dorsal root ganglia (DRG). This underscores the fact that both peripheral ganglia consist of diverse and distinct populations of cells best suited for their respective functions. In this regard, postganglionic neurons, akin to motor neurons, primarily control smooth muscle contractions like that of blood vessels by releasing NE at the synapse, unlike their sensory counterparts. While the human DRG and trigeminal ganglia have recently been characterized using spatial and single nuclei RNA sequencing methods (10-15), the human SCGs have not been molecularly characterized. This creates a gap in knowledge that impacts drug development. For instance, recent efforts to develop analgesics for chronic pain such as Nav1.7 inhibitors, have been discontinued due to severe sympathetic side effects like bradycardia and hypotension (16-19). This observation is substantiated by prior work assessing Nav1.7 expression and currents in sympathetic neurons which are impacted by gain-of-function mutations associated with erythromelalgia and small-fiber neuropathy (20-23).

Hence, understanding the molecular characterization of cells in SCGs presents a unique opportunity for selectively targeting the sympathetic nervous system or preserving sympathetic function when targeting the sensory nervous system.

We hypothesized that human SCGs would consist of unique populations of neuronal and non-neuronal cells, with diabetes inducing transcriptomic changes in these populations. To test this, we used spatial transcriptomics of human lumbar SCGs recovered from organ donors to identify 3 distinct populations of sympathetic neurons, and 9 clusters of non-neuronal cells. In diabetic samples, we noted a significant decrease in a group of myelinating Schwann cells and a reprogramming of CHO neurons, likely due to an imbalance of target-derived neurotrophic support. By comparing lumbar SCG and DRG neurons, we identified a unique set of genes specific to each ganglion. Our data gives unprecedented insight into the molecular and cellular complexity of the human paravertebral SCG.

## Results

### Characterization of the lumbar sympathetic chain ganglia (SCG)

Lumbar SCGs were recovered from 6 organ donors (3 males, 3 females) with and without a history of type 2 diabetes mellitus (**Sup. Table 1**). The SCGs were identified as a string of ganglia attached laterally to the ventral spinal column. The ventral vertebrae were removed as described previously during DRG recovery (**Fig. 1A, B**). The SCGs were immediately cleaned of fat and connective tissue and fresh frozen in pulverized dry ice for subsequent analysis. By performing hematoxylin and eosin (H&E) staining, we confirmed the presence of neuronal cell bodies – replete with lipofuscin – axon bundles, non-neuronal cells, and the epineurium (**Fig. 1C**). Notably, human SCG neurons were compactly organized, unlike the relatively more diffuse neuronal distribution of the human DRG previously reported (24).

**Table 1.**
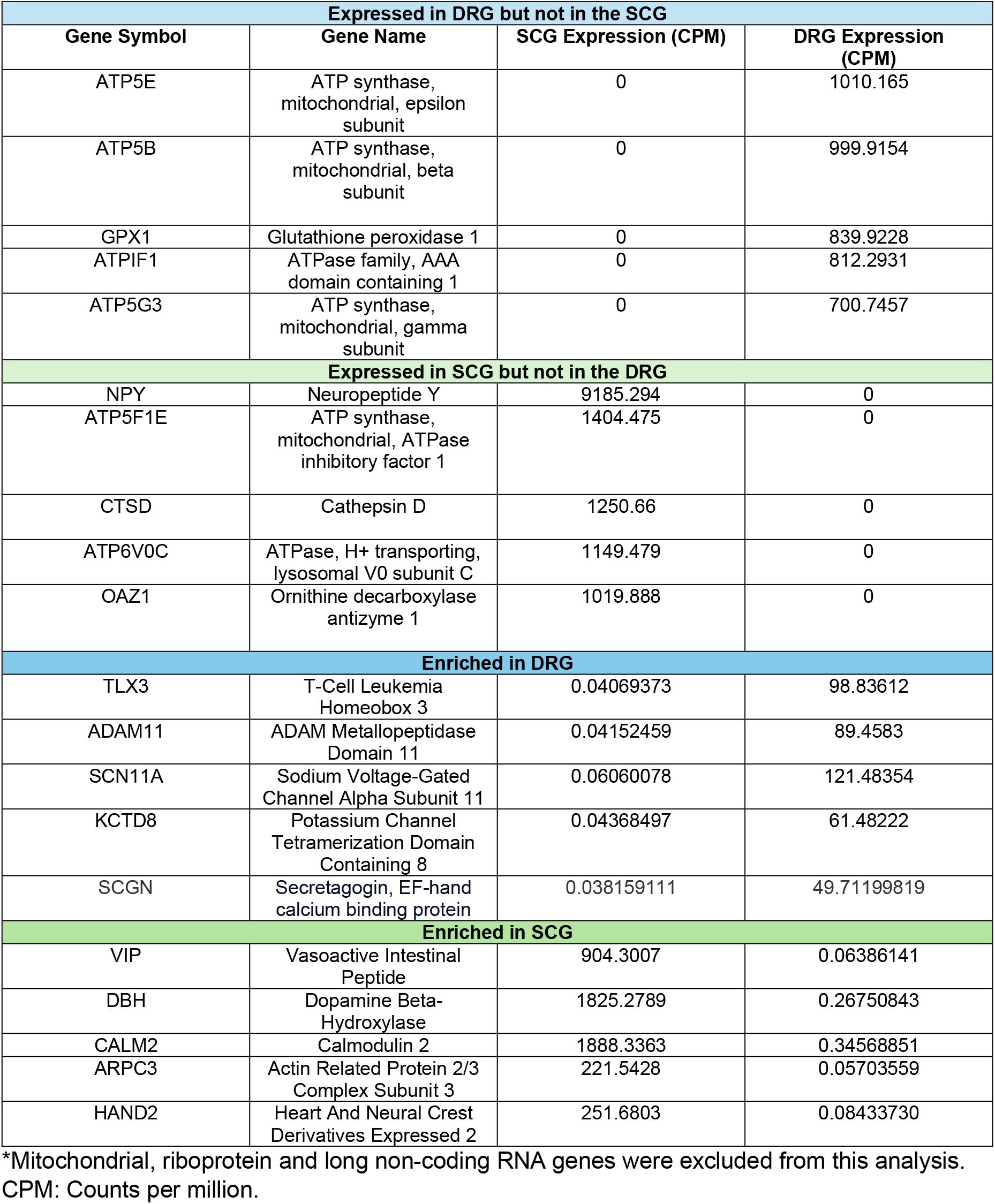
Top 5 genes expressed exclusively or preferentially in the DRG and SCG. *Mitochondrial, riboprotein and long non-coding RNA genes were excluded from this analysis. CPM: Counts per million.

**Figure 1.**
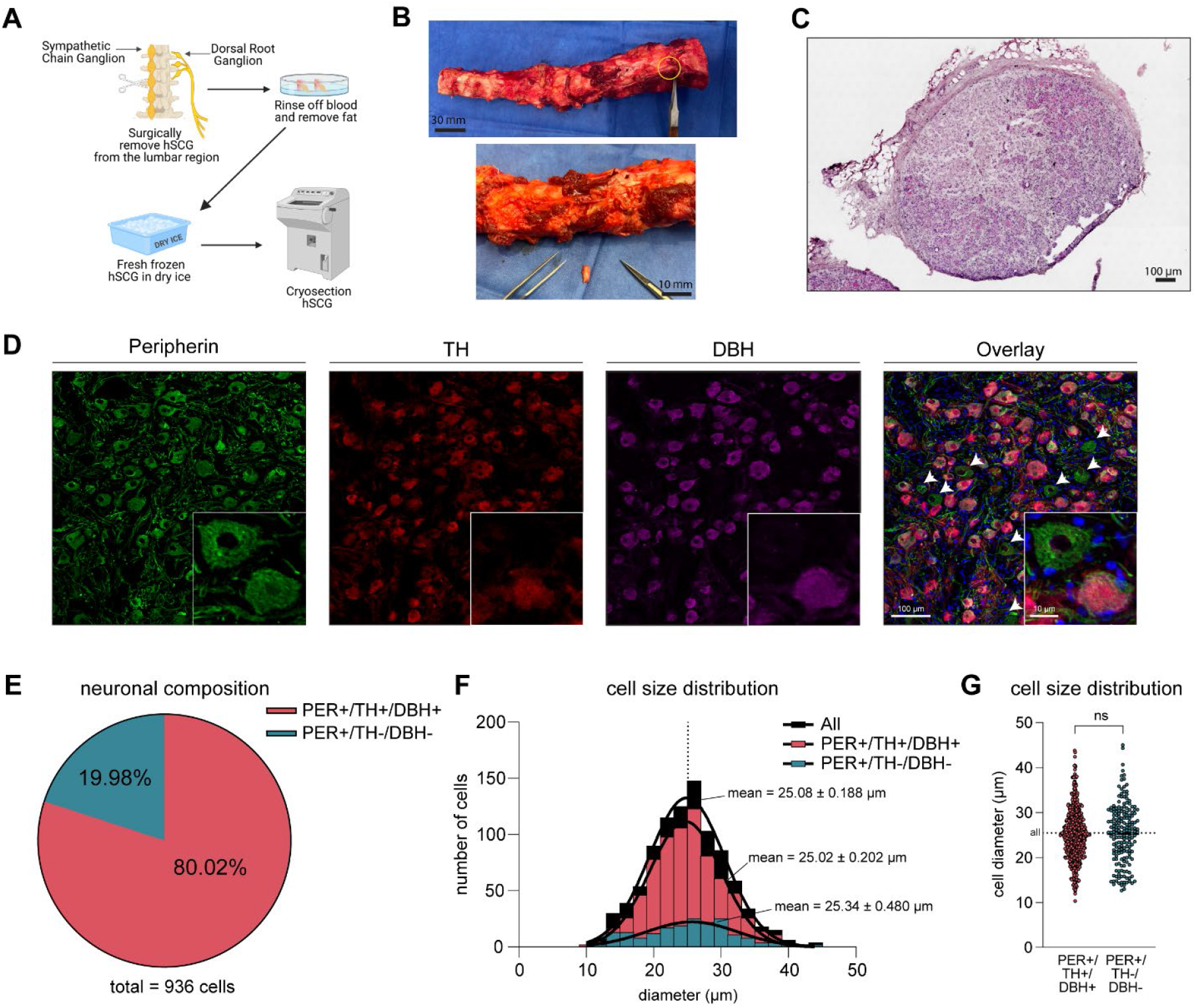
Sympathetic Chain Ganglia (SCG) surgical recovery and quality control assessment. A, B. Human SCG, isolated from the lumbar region of the vertebral column, was surgically harvested from organ donors. The tissue was rinsed with artificial cerebrospinal fluid (aCSF) to remove blood, followed by freezing in powdered dry ice and cryosectioning. **C**. Tissue quality was evaluated by Hematoxylin and Eosin (H&E) staining. Cell bodies and fibrous tissue appear rose-colored, while the nuclei are stained deep blue-purple. No freezing artifacts were observed. **D, E**. Immunohistochemical analysis revealed that 80.02 % were PER+/TH+/DBH+, while 19.98% were PER+/TH-/DBH-(Peripherin = green, TH = red, and DBH = purple). **F, G**. Neuronal size distribution and composition were quantified, with sizes ranging from 10 μm to 45 μm. No significant differences were observed in the size distribution between both populations (two-tailed unpaired t-test). Data presented as mean ± standard error of mean (SEM). NS, not significant.

Tyrosine hydroxylase (TH) and dopamine β-hydroxylase (DBH) are key proteins in the synthesis of norepinephrine and serve as markers of noradrenergic neurons of the autonomic nervous system (25). We performed immunohistochemistry of non-diabetic SCGs (n=2 donors, 3 sections, 100 µm apart, per donor) and identified that the majority (∼80%) of all SCG neurons were TH- and DBH-positive, indicative of a noradrenergic phenotype (**Fig. 1D, E**). The remaining ∼20% of neurons were immunoreactive for peripherin (*PRPH*), a cytoskeletal protein specifically expressed in neurons, but negative for TH and DBH. This suggests that these PER+/TH-/DBH-were putative cholinergic neurons, consistent with prior reports of human and rodent sympathetic ganglia (9). We did not find any neuron that expressed only TH or DBH. In the rat inferior mesenteric ganglion, cholinergic neurons were found to be larger than noradrenergic neurons (5). Thus, we next examined the cell size of TH+/DBH+ and TH-/DBH-neurons in the human SCG and found no significant difference between groups (**Fig. 1F, G**). Interestingly, the overall size of SCG neurons was consistent among mice, rats, guinea pigs, and humans across a narrow size distribution profile unlike DRG neurons (5, 26, 27).

### Sympathetic neurons of the SCG

To investigate the neuronal composition of the SCG, we used the 10X Genomics Visium spatial sequencing technology combined with computational analysis using the Seurat graphed-based workflow (see Methods) (**Fig. 2A**) (**Sup. Table 2**) (28). By generating a list of differentially expressed genes (DEGs) per cluster (**Sup. Table 3**), we identified three distinct neuronal populations: two noradrenergic (NA1 and NA2) and one cholinergic (CHO) (**Fig. 2B**), all spatially localized within the core region of the SCG (**Fig. 2C, Sup. Fig. S1**). We were unable to visually identify any specific distributive patterns among the neuronal populations. Both noradrenergic populations (NA1 and NA2) expressed transcripts for *TH, DBH, ARHGAP36* (Rho GTPase-activating protein 36), and *SLC6A2* (solute carrier family 6 member 2, also known as norepinephrine transporter NET). NA1 and NA2 populations were discriminated by lower expression of peripherin (*PRPH*), N-Myc downstream regulated gene 4 (*NDRG4*), and secretogranin 2 (*SCG2*) in NA1. The cholinergic population was characterized by minimal *TH* expression but was defined by the expression of *VIP, SLC18A3* (solute carrier family 18 member A3 or vesicular acetylcholine transporter VAChT), and *CALCA* (calcitonin-related polypeptide alpha) (8) (**Fig. 2D–H**). We were unable to identify a nitrergic population based on *NOS1* (nitric oxide synthase 1) expression, unlike a previous report in rat stellate ganglia, which are part of the cervicothoracic level of the SCGs. We compared the gene expression profiles across the three neuronal populations to assess their similarities. A total of 65 genes were shared among all three populations (**Sup. Table 4**). As expected, the largest overlap was observed between two noradrenergic populations (NA1 and NA2), with 1073 genes common to both populations. In comparison, smaller overlaps between NA1 and CHO (60 genes) and NA2 and CHO (127 genes) were identified. Furthermore, 338 genes were uniquely expressed in NA1, and 224 genes in NA2. CHO exhibited the least complexity with only 64 genes expressed exclusively within this population (**Fig. 2I**).

**Figure 2.**
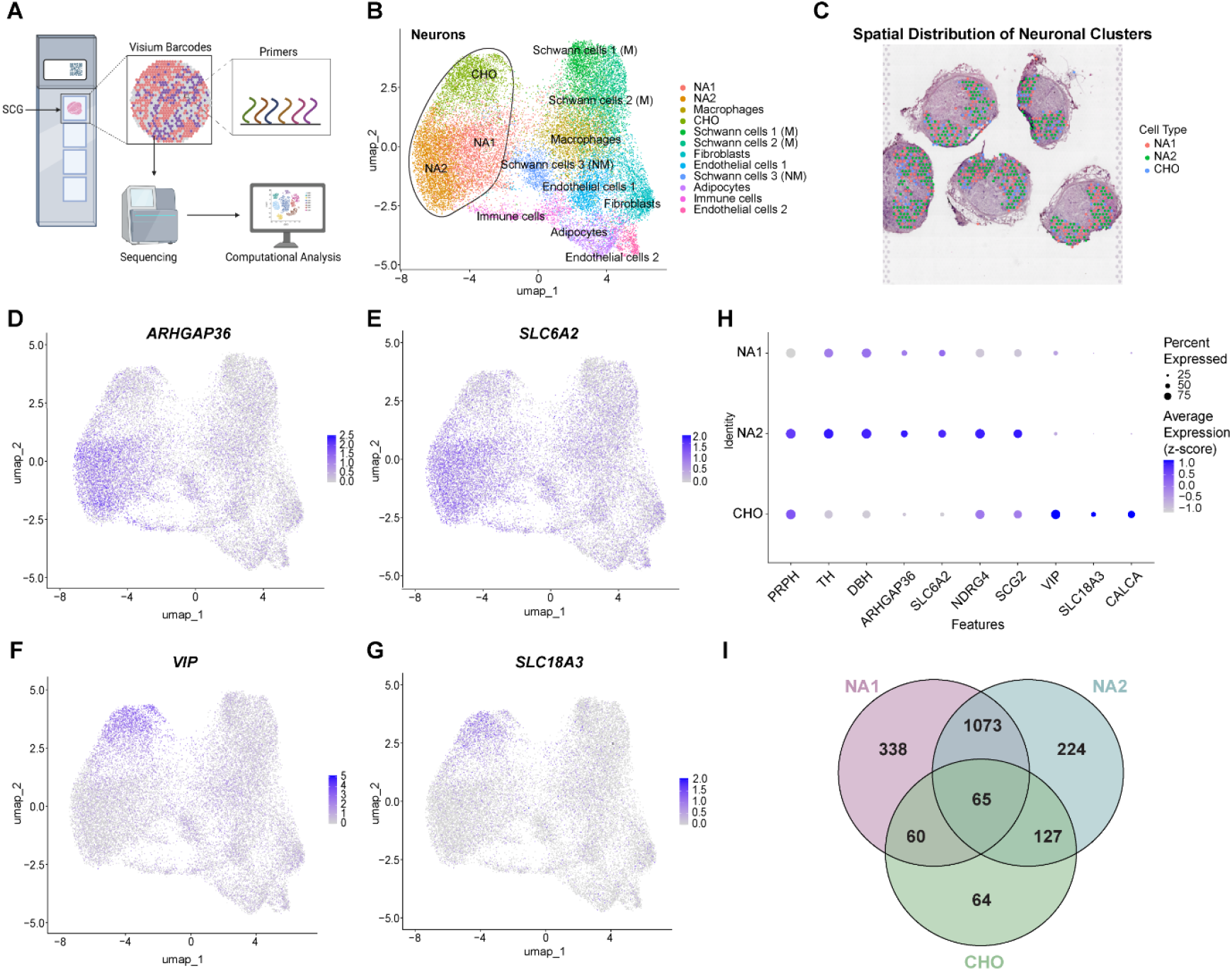
Identification of neuronal populations in the SCG. **A**. Workflow for spatial sequencing of human SCG tissue. Fresh-frozen SCG sections were mounted on 10X Genomics Visium slides within the fiducial frames. After permeabilization, mRNA hybridization was performed using primers on the Visium barcoded spots. The samples were then sequenced and analyzed using the Seurat package in R. **B**. UMAP plot of neuronal clusters in the SCG, revealing three distinct clusters: NA1, NA2, and CHO. **C**. Spatial distribution of neuronal clusters within SCG tissue sections. Colored overlapping barcodes indicate each neuronal population, which were all localized in the core of the SCG. **D, E**. Feature plots for *ARHGAP36* and *SLC6A2*, markers of noradrenergic neurons, showing highly localized expression in the NA1 and NA2 clusters. **F, G**. Feature plots for *VIP* and *SLC18A3*, markers of cholinergic neurons, demonstrating high expression within the CHO cluster. **H**. Dot plot showing expression of select genes across neuronal populations. The circle size represents the percentage of cells expressing each gene, and the grey-to-blue gradient indicates increasing average gene expression. NA1 shows low expression of peripherin (*PRPH*), *NDRG4* and *SCG2*. Both NA1 and NA2 express *TH, DBH, ARHGAP36*, and *SLC6A2*, while CHO expresses *VIP, SLC18A3*, and *CALCA* at high levels. **I**. Venn diagram illustrating gene expression overlap across neuronal populations. Intersections reveal 65 genes common to all three populations. Additionally, NA1, NA2, and CHO clusters expressed 338, 224, and 64 unique genes, respectively.

### Non-neuronal cells of the SCG

We further explored the composition of non-neuronal cells of the SCG. Our analysis identified nine distinct non-neuronal populations, including adipocytes, endothelial cells (two subpopulations), fibroblasts, immune cells, macrophages, and Schwann cells (three subpopulations, 2 myelinating and 1 non-myelinating) (**Fig. 3A, Sup. Fig. S2**). The majority of these cells were spatially localized surrounding the neuronal core of the SCG. (**Fig. 3B**). Using Seurat, we identified differentially expressed genes (DEGs), which were used to distinguish each cell type using the EnrichR platform (**Fig. 3C**). Through this approach we identified an adipocyte cluster characterized by the expression of adiponectin (*ADIPOQ*), and fatty acid-binding protein 4 (*FABP4*). We also identified two endothelial cell populations, both expressing aquaporin 1 (*AQP1*), endoglin (*ENG*), platelet endothelial cell adhesion molecule 1 (*PECAM1*), and claudin 5 (*CLDN5*). The second endothelial subset exhibited additional expression of complement factor D (*CFD*), myosin heavy chain 11 (*MYH11*), actin alpha 2 (*ACTA2*), and cysteine-rich protein 1 (*CRIP1*), suggesting a more specialized vascular function. Fibroblasts were characterized by high expression of *CRIP1*, matrix Gla protein (*MGP*) and glutathione peroxidase 3 (*GPX3*). The immune cell population displayed a transcriptional signature consistent with B cells, marked by high expression of CD34 molecule (*CD34*), and immunoglobulin components (*IGKC, IGHG1, IGHA1, IGLC2*, and *IGHG3*). Macrophages expressed a combination of *MGP* and *GPX3*. Notably, we identified three distinct Schwann cell clusters. Schwann cell populations 1 and 2 were classified as myelinating Schwann cells due to their high expression of myelin protein zero (*MPZ*), with the second population distinguished by additional elevated expression of periaxin (*PRX*). The third Schwann cell population was classified as non-myelinating Schwann cells, characterized by expression of S100 calcium-binding protein (*S100B*) and a high expression of mitochondrial *MTRNR2L12* gene. We were unable to confidently cluster satellite glial cells using this approach, due to the close proximity of satellite glial cells to neurons and limitations of the spatial transcriptomics technique as a near-single cell RNAseq approach. Nonetheless, we identified and defined 9 non-neuronal cell populations that constitute the human SCG (**Sup. Table 3**).

**Figure 3.**
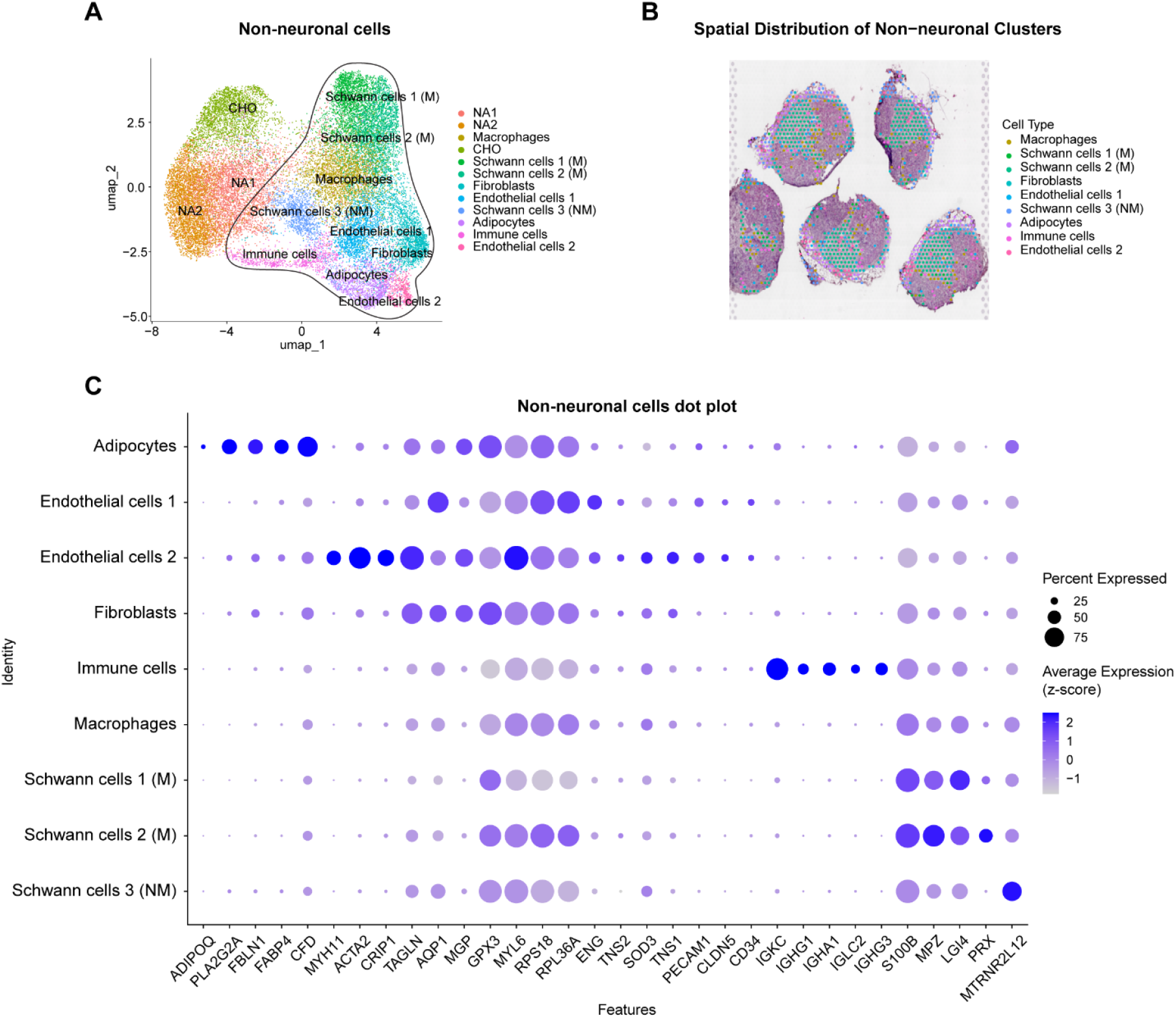
Identification of non-neuronal populations in the SCG. **A**. UMAP plot of spatial transcriptomics data from human SCG tissues reveals nine distinct non-neuronal cell populations. **B**. Spatial distribution of these non-neuronal clusters within tissue sections, where overlapping colored barcodes highlight specific non-neuronal cell types particularly localized surrounding the neuronal core of the SCG. **C**. Dot plot illustrating gene expression profiles in the non-neuronal cell types. Schwann cells were classified into two populations, myelinating (M) and non-myelinating (NM). The circle size represents the proportion of cells expressing each gene, while the white-to-blue gradient indicates increasing average gene expression levels. The x-axis shows specific genes, and the y-axis corresponds to non-neuronal cell types.

### The Effect of Diabetes on SCGs

Anatomic and cellular changes in the sympathetic ganglia of diabetic patients have been recognized since the mid-1960s (1, 29). We performed spatial transcriptomics on SCGs from 2 diabetic and 4 non-diabetic (ND) organ donors to investigate the transcriptomic changes induced by diabetes. We obtained a total of 10,709 neurons, with 7,385 cells derived from non-diabetic samples and 3,324 neurons from diabetic samples (**Sup. Table 5**). We first sought to compare the proportions of each cell type from each condition. We found that the SCG consisted of 39.9% and 41.8% of neurons in the ND and diabetic SCGs, respectively, confirming that our diabetic samples did not suffer from neuronal loss (**Fig. 4A**), unlike in the DRG where significant neuronal loss is observed (30). We then assessed the distribution of subtypes within the larger neuronal population. We found that the distribution of NA and CHO neurons was skewed in the diabetic condition, with a lower proportion of CHO cells (27.6% vs 18%) and a higher proportion of NA cells (38.7% vs 44.6% NA1, and 33.7% vs 37.4% NA2) (**Fig. 4B**). To statistically test shifts in cell proportions, we employed the Policastro test (FDR<0.05, Log2 fold difference>0.5) and found a significant decrease in a myelinating Schwann cell population (Schwann cells 1 (M)) and CHO populations in the diabetic condition, while endothelial cell 1 population showed a significant increase (**Fig. 4C**) (**Sup. Table 6**).

**Figure 4.**
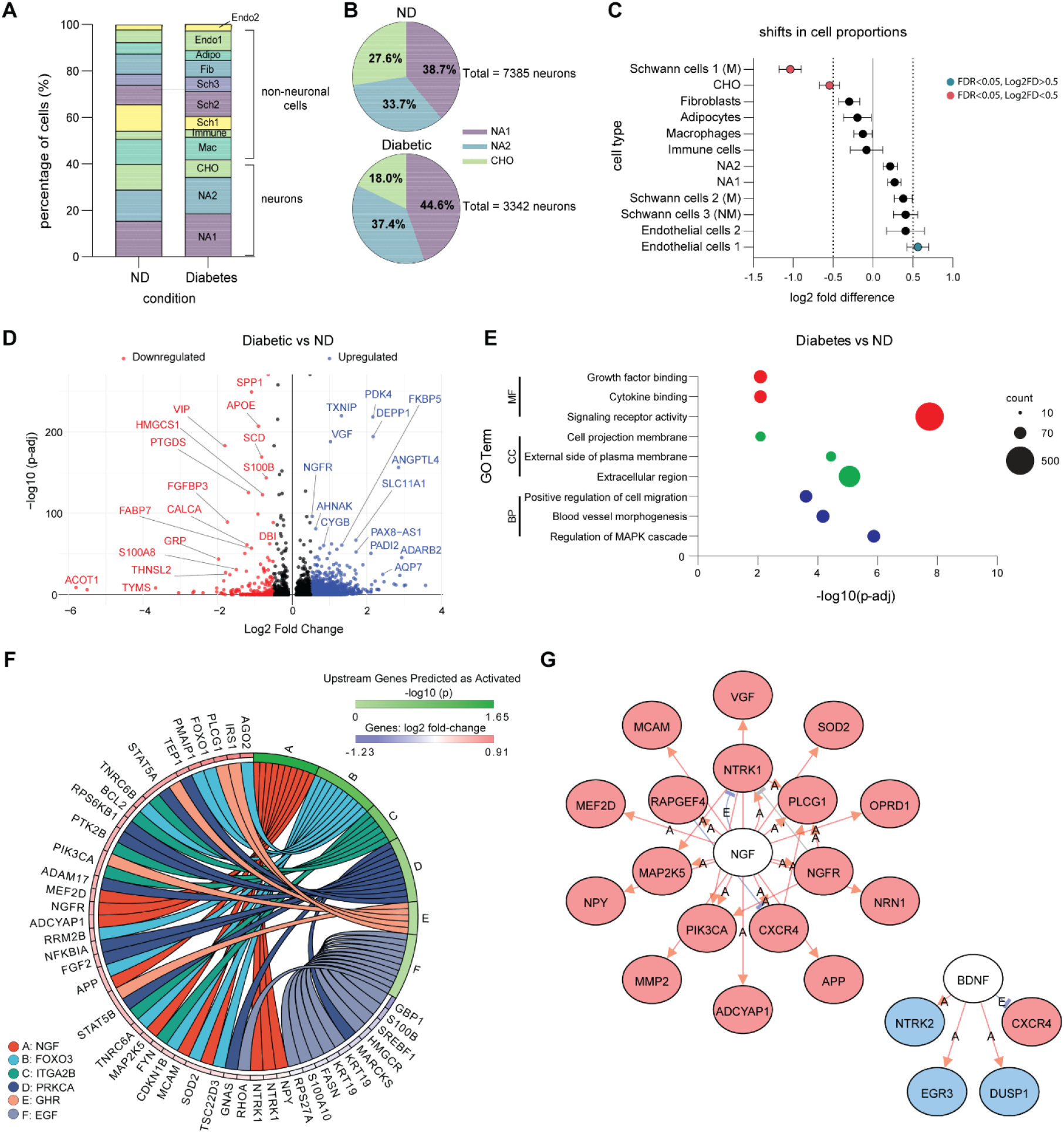
Diabetes alters SCG cell composition and gene expression. **A**. Distribution of cell populations in non-diabetic (ND) and diabetic conditions, illustrating a shift in cellular composition. **B**. Proportions of neuronal subpopulations in ND (7,385 neurons) and diabetic (3,324 neurons) conditions show a shift in population composition: 38.7% NA1, 33.7% NA2, and 27.6% CHO in ND, versus 44.6% NA1, 37.4% NA2, and 18.0% CHO in diabetic conditions. **C**. Differential cell proportion analysis confirms a significant decrease in Schwann cell 1 (M) and CHO populations in the diabetic condition, while endothelial cells 1 increase (Policastro test p < 0.05 & Log2F > 0.5). **D**. Volcano plot of differentially expressed genes shows significant upregulation (blue) and downregulation (red) of genes in the diabetic condition. **E**. Gene Ontology (GO) enrichment analysis of differentially expressed genes identifies affected biological processes, cellular components, and molecular functions in diabetic SCG neurons. Circle size represents the number of genes in each category (10-500). **F**. Circos plot showing the interactions of differentially expressed genes across various signaling pathways impacted by diabetes. **G**. Gene interaction network highlights two main clusters: one centered around NGF (nerve growth factor) and the other around BDNF (brain-derived neurotrophic factor). Red indicates a significantly upregulated gene, while blue indicates a significantly downregulated gene.

For the next set of analyses, we focused on the neurons and analyzed differentially expressed genes in the diabetic condition using Seurat. We identified 276 downregulated, and 1972 upregulated DEGs. *VIP* and *CALCA*, genes specific to the CHO population, were among the downregulated genes, while *DEPP1*, an insulin-regulatory protein, was upregulated (**Fig. 4D**, see **Sup. Table 7** for a detailed list). To increase the reliability of our findings, we confirmed a reduction in *VIP* and an increase in *DEPP1* mRNA using qPCR in a separate cohort of SCGs from diabetic and ND donors (**Sup. Fig. S3**). This A Gene Ontology (GO) pathway analysis was done to identify molecular functions (MF), cellular components (CC), and biological processes (BP) associated with the DEGs (**Sup. Tables 8-10**). For MF we observed an enrichment in signaling receptor activity, growth factor binding and cytokine binding. For CC, the most enriched categories included the extracellular region, external side of plasma membrane, and cell projection membrane. Lastly, for BP the DEG were primarily involved in regulation of MAPK cascade, blood vessel morphogenesis, and positive regulation of cell migration (**Fig. 4E**). We identified several key factors and signaling molecules, particularly *NGF, FOXO3, ITGA2B, GHR*, and *EGF*, as activated upstream regulators in the diabetic SCG (**Fig. 4F**). Further analysis revealed notable differential regulation of neurotrophic factor pathways. Specifically, NGF-related pathways were upregulated, with genes such as *NTRK1, NGFR*, and *MAP2K5*. In contrast, BDNF-dependent pathways exhibited downregulated expression of genes, including *NTRK2, EGR3* and *DUSP1* (**Fig. 4G**) (**Sup. Table 11**). Taken together, these data suggest that diabetes is associated with an imbalance in potentially target-derived neurotrophic support and ultimately a change in neuronal phenotype from cholinergic to noradrenergic.

### Interactions between sensory and sympathetic neurons

Sympathetic neurons of the SCG control blood flow to the DRG and in painful pathological conditions in rodent models, SCG neurons can form pericellular nests around DRG neurons to modulate the activity of nociceptors (31). To explore the potential interactions between sympathetic and sensory neurons, we performed an interactome analysis using our dataset and a previously published spatial transcriptomics dataset of DRG neurons (**Sup. Fig. S4**) (11, 32). We tested two possibilities: (1) an SCG ligand that acts on a DRG neuron receptor to alter its excitability, (2) a DRG ligand that acts on an SCG receptor to promote sympathetic outgrowth, innervation, and the formation of pericellular nests. Our analysis identified eight SCG ligands interacting with 48 DRG receptors, with neuropeptide Y (*NPY*) and clusterin (*CLU*) showing the most interactions. Additionally, we found 46 DRG ligands, including receptors for products of the phospholipase A2 (*PLA2*) pathway and Leucine Rich Repeat and Fibronectin Type III Domain Containing (*LRFN*) families, interacting with 18 SCG receptors, such *NGFR* (**Sup. Fig. S3, Sup. Table 12**). These data present possible ligand-receptor interactions that give rise to aberrant outgrowth of sympathetic fibers in the DRG, a potential cause of chronic neuropathic pain .

### Distinct gene sets in sympathetic and sensory neurons

We next sought to compare the expression of well-characterized analgesic targets in the DRG and SCG, focusing on sodium channels, TRP channels, potassium channels, and opioid receptors (**Fig. 5**) (33, 34) . A significant finding was the absence of *SCN10A* (encoding Nav1.8) expression in the SCG, confirming with a lack of sympathetic side effects of suzetrigine, a recently approved Nav1.8 inhibitor for acute pain conditions (35). Additionally, genes such as *KCNC2*, and *KCNK9* were not expressed in the GSCG. Unlike DRG neurons, *SCN1A* (Nav1.1) and *SCN11A* (Nav1.9) were minimally expressed in SCG neurons. We further conducted a presence-absence and enrichment analysis of all genes expressed in the neurons of both tissue types, identifying genes that were uniquely expressed or enriched in either SCG or DRG neurons. The top 5 genes per tissue are indicated in **Table 1** and a complete list is compiled in the **Sup. Tables 12-14**. Notably, glutathione peroxidase 1 (*GPX1*), a target of interest for diabetic neuropathy induced by oxidative stress, was exclusively expressed in DRG neurons (36, 37). These data highlight important transcriptomic differences between neurons of the SCG and the DRG that are relevant drug development efforts.

**Figure 5.**
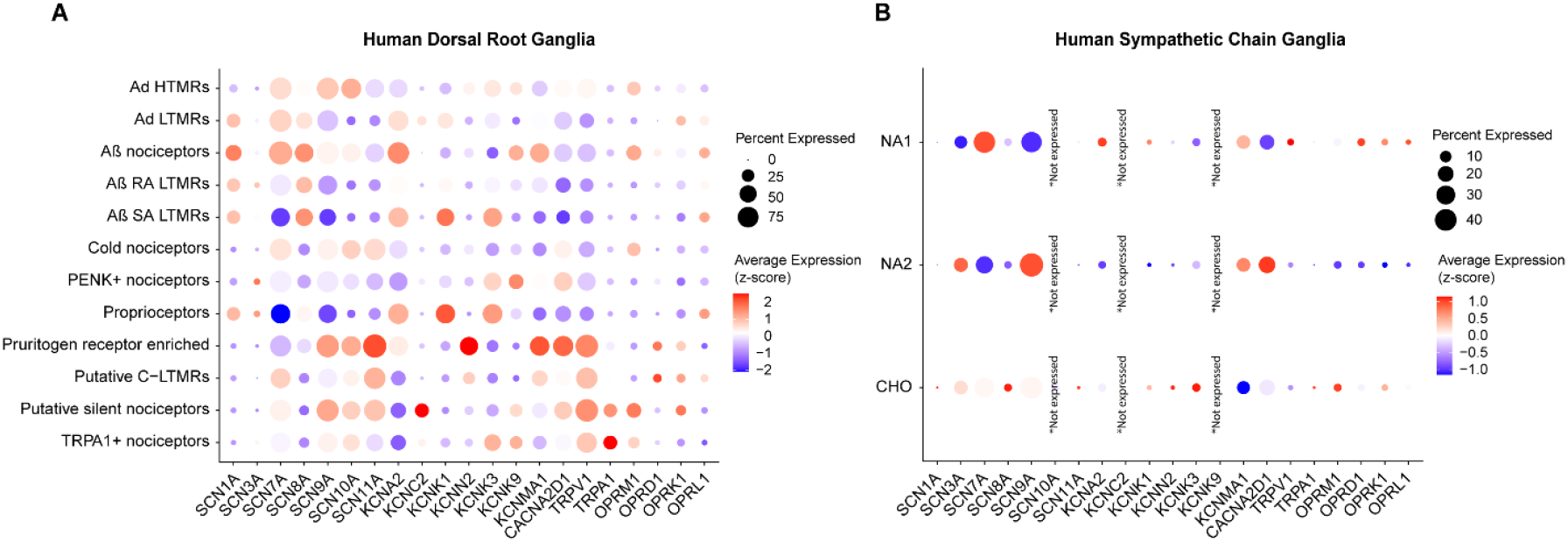
Gene expression comparisons between sensory and sympathetic neurons. Dot plots showing differential expression of well-known gene targets for drug development including sodium-potassium channels, TRP channels, and opioid receptors across **A**. DRG and **B**. SCG, where *SCN10A, KCNC2* and *KCNK9* were not expressed. Circle size represents the percentage of cells expressing each gene. The average expression is shown on a blue-to-red gradient, where blue indicates the lowest expression and red the highest.

## Discussion

Previous studies characterizing the human sympathetic ganglia were constrained by biased approaches and low-throughput techniques, such as immunohistochemistry and in situ hybridization, which lacked the depth, resolution, and scalability of high-throughput methods like RNA sequencing. We aimed to characterize the cellular makeup of lumbar sympathetic chain ganglia (SCG) and identify molecular mechanisms underlying sympathetic dysfunction in diabetes, ultimately highlighting potential therapeutic targets. Using spatial transcriptomics, we identified 12 distinct populations of cells in the lumbar SCG: 3 neuronal and 9 non-neuronal cell subtypes. We discovered a partial phenotypic switch of CHO neurons in the diabetic SCGs when compared to non-diabetic SCGs. Further investigation also revealed reduced mRNA expression of neuropeptides like *VIP*, and *CALCA* in CHO neurons. By comparing the lumbar SCG to lumbar DRGs, we identified a distinct set of genes expressed in both ganglia creating a compendium of possible druggable targets.

Over the past decade, RNA sequencing technologies have led to an unbiased characterization of rodent SCGs (8, 9). Using single cell RNA sequencing, Furlan, *et al*. (8) describe neurons of the thoracic SCG from mice in detail, identifying 7 neuronal populations: 5 noradrenergic and 2 cholinergic. Subsequent work analyzed non-neuronal cells of the mouse SCG and discovered a unique transcriptomic profile associated with sympathetic satellite glial cells. Moving rostrally, Sharma, *et al*. (38) identified multiple noradrenergic and cholinergic neuronal populations in the mouse stellate ganglia. Using viral tracers, they identified 3 noradrenergic populations that specifically projected to the heart and not the paw. More recently, prevertebral ganglia of the mouse, particularly the celiac and superior mesenteric ganglia, have been sequenced (2, 39). While these studies provide unprecedented insight into the composition and activity of sympathetic ganglia, they are limited by species differences, particularly considering the number of recent studies showing divergence in the composition of mouse and human DRGs I(14, 27, 40). In our human dataset, we were unable to find a number of neuronal classification genes corresponding to murine transcription factors (*Tlx3, Tbx20, Barx2*), receptors (*Rxfp1, Adrbk2, Npy4r, Sctr, Cckar, Ltk, Gpr1, Cckbr, Agtr2*), and neuropeptides (*Tac2, Nppc, Cbln2*). While not directly tested in this paper, the transcriptomic signatures of autonomic neurons are likely to differ between mice and humans, an area that deserves further attention in the future.

An earlier report found that 75% of human thoracic SCG neurons express both TH and DBH mRNA transcripts (41). Similarly, we observed that 80% of neurons in the lumbar SCG express TH and DBH proteins (**Fig. 1**). This contrasts with the superior cervical, middle cervical, and stellate ganglia, where over 90% of neurons are DBH-positive (42) supporting the conclusion that the molecular composition of SCGs varies by spinal level. Previous work has suggested that innervation targets shape the molecular identity of neurons within the sympathetic nervous system (3). This raises the intriguing possibility that diseases or injuries affecting innervated targets could induce persistent molecular changes in sympathetic neurons, potentially leading to sympathetic complications secondary to the disease.

Consistent with this, we found an altered CHO neuron phenotype within the SCG in diabetes, which may contribute to autonomic dysfunction and altered sympathetic outflow. Cholinergic neurons in the SCG originate from noradrenergic precursors during development, undergoing a phenotypic shift driven by target-derived signals and transcriptional programs (43). Interestingly, injured neurons in the DRG can reactivate developmental pathways to promote survival and regeneration (44, 45). Given this plasticity, it is not a surprise that our findings suggest that cholinergic neurons in the SCG may undergo a phenotypic switch back to a noradrenergic state in response to a diabetic pathology. While what exactly causes this phenotypic switch remains to be discovered, our RNA sequencing dataset provides clues for developing testable hypotheses. A key finding from our comparison of diabetic and non-diabetic sympathetic neurons was a shift in neurotrophic signaling, particularly an increase in NGF and a decrease in BDNF expression. A few lines of evidence support this hypothesis. First, NTRK1 and NTRK2 are specifically expressed in noradrenergic and cholinergic neurons, respectively, suggesting that these two receptors are key to the identity of each cell type (8, 9). Second, several studies have demonstrated that NGF/TRKA is critical for the differentiation and maintenance of adrenergic neurons (46, 47). Similarly, BDNF/TRKB signaling in sweat glands, the primary target of cholinergic sympathetic neurons, is an embryonic driver of cholinergic switch from an adrenergic precursor (48, 49). There could be two consequences of such a change: (1) cholinergic neurons regulate vasodilation and thermoregulation, and hence the loss of these cells can impair blood flow and sweating, a common phenomenon in diabetes; (2) a phenotypic shift can increase noradrenergic tone, a known contributor to diabetic heart disease, hypertension, and microvascular dysfunction (50). Compounding the loss of cholinergic phenotype is a reduction in transcripts for vasodilator neuropeptides like *VIP* and *CALCA* (encoding CGRP) in the diabetic SCG. By characterizing the cholinergic population of the human SCG, our data lays the foundation for precise, testable hypotheses that could uncover the molecular mechanisms governing this phenotypic switch, ultimately guiding the development of targeted therapeutic interventions.

Understanding the transcriptome of both sympathetic and sensory ganglia is critical for developing targeted therapeutics that selectively modulate each system while minimizing unintended effects on the other. A key example of this is the development of Nav1.7-targeted analgesics for neuropathic pain (51-53). While Nav1.7 is a well-established target in sensory neurons, its expression in sympathetic ganglia led to unexpected off-target effects on autonomic function, including dose-dependent decreases in heart-rate variability and syncope encountered during clinical development and recapitulated in non-human primates (17-19). These findings are consistent with our observation here that *SCN9A* is the most highly expressed voltage gated sodium channel in human SCG and a separate report that a loss-of-function mutation of Nav1.7 contributes to autonomic dysfunction (54). In contrast, Nav1.8 antagonists, which are effective for acute pain relief, do not produce these autonomic side effects because *SCN10A*, the gene encoding Nav1.8, is not expressed in sympathetic neurons. Similarly, *SCN11A* (Nav1.9) expression is limited in the SCG neurons suggesting that targeting Nav1.9 could be a therapeutic strategy for pain without autonomic issues. Nav1.3 (encoded by *SCN3A*) serves as a sympathetically-enriched target, as it is minimally expressed in human DRGs (11, 13, 14) but is one of the predominant sodium channels in SCG neurons (**Fig 5**). This data highlights the importance of transcriptomic profiling in distinguishing targets that are exclusive to either the sensory or sympathetic nervous system, thereby enabling the development of safer, more precise therapeutics.

We acknowledge that there are some limitations to our findings. Firstly, the spatial transcriptomics technique provides valuable spatial context for gene expression at a *near*-single-cell resolution. While this approach could potentially lead to signal averaging across multiple cell types within a given region, our data suggests that spatial transcriptomics sufficiently capture multiple neuronal and non-neuronal populations known to exist in sympathetic ganglia. An exception to this is the transcriptomic profile of satellite glial cells which are overshadowed by the neurons which they surround. Secondly, our analysis is constraint by a low number of donor samples which may limit the generalizability of our findings regarding diabetes-associated transcriptional changes. However, the robust differences between diabetic and non-diabetic samples obtained in this report provide a foundation for more specific hypotheses that can be tested in the future. Thirdly, the lack of detailed medical history from organ donors prevents a comprehensive understanding of comorbidities or prior treatments that could influence gene expression patterns, potentially introducing an unknown layer of variability in our dataset. Finally, while our data robustly explored multiple hypotheses related to sympathetic dysfunction in diabetes, we were unable to confirm mechanistic causality in this manuscript. The data presented in this paper, however, implicate certain cell types, CHO neurons and myelinating Schwann cells, and mechanisms, such as altered neurotrophic support and vasodilatory neuropeptides in diabetes, that can be explored further.

## Materials and Methods

### Tissue Recovery

Human lumbar L1-L2 (SCG) were recovered in collaboration with the Southwest Transplant Alliance, following a previous publication (55). Donor clinical history and further details are elaborated in Supplemental Materials (**Sup. Table 1**). The study was approved by the Institutional Review Board at the University of Texas at Dallas.

### Immunohistochemistry & Imaging

We followed a modified immunocytochemistry protocol based on Yousuf, *et al*. (56).

Details of this experiment can be found in the **supplementary methods**.

Imaging was performed using an FV3000 confocal microscope (Evident Scientific) at 20x magnification, with brightness and contrast adjustments made in Olympus CellSens software (v1.18).

### 10X genomics Visium spatial transcriptomics

Tissue preparation, fixation, staining, and imaging was conducted exactly as described by and 10X *Genomics (28)*. The optimal permeabilization time for human SCG RNA release was determined to be 30 minutes. Full results are in **supplementary methods**. The Visium spatial gene expression library and sequencing were conducted at the Genomics Core facilities of the University of Texas at Dallas. The samples underwent paired-end sequencing using the 10X Genomics Visium library and the Illumina NextSeq 2000 sequencing system. Sequencing details can be found in supplementary materials.

### Visium Spatial RNA-seq Analysis

We processed Visium data using the 10X Genomics Space Ranger pipeline (v1.1). Illumina BCL files were processed using spaceranger mkfastq to generate FASTQ files, which were then used by spaceranger count along with bright-field images and the human reference transcriptome (GRCh38.p10). The resulting output folders, containing tissue images, coordinates, and barcode and gene expression counts, were analyzed using the Seurat package (v5.0.2) in R (v4.3.3) following the “Analysis, visualization and integration of spatial datasets with Seurat” pipeline developed by the Satjia Lab (57). Data were normalized, and cell clusters were identified using FindNeighbors (dims = 1:20), FindClusters (resolution = 1), and RunUMAP (dims = 1:20). Batch correction for all barcodes was performed with the Harmony algorithm (dims = 1:20) (58). Differentially expressed genes were identified using the Wilcoxon rank-sum test in Seurat, and the Policastro test assessed cell proportion differences between clusters, calculating p-values<0.05 via permutation testing and confidence intervals through bootstrapping.

### Differentially Expressed Genes in Diabetic Conditions

We generated a list of Differentially Expressed Genes per condition (Non-diabetic vs. Diabetic) using the FindAllMarkers() function on subset neuronal clusters (NA1, N2, CHO) with a LogFold threshold of 0.25, p.value < 0.05 and test = “Wilcox”.

### Gene Ontology (GO) analysis

We uploaded the previously generated DEG list from neuronal clusters per condition as a .txt file into Advaita’s next-gen pathway analysis tool, “iPathway”. The analysis was performed using the *Homo sapiens* reference, including all genes in the alignment, with a significance threshold of p < 0.05 and a log-fold change cutoff of 0.5.

See the Methods section in the supplementary materials for more details.

## Data availability

Datasets generated from this study will be publicly available through https://sensoryomics.shinyapps.io/RNA-Data/ and the SPARC portal.

*Figures were generated in R, BioRender, GraphPad Prism 10 and Adobe Illustrator 2024*.

## Acknowledgments and funding sources

We are grateful to the organ donors and their families for the gift of life and research provided by their organ donation.

