## Supplementary material for "Spatial transcriptomic profiling of human paravertebral sympathetic chain ganglia reveals diabetes-induced neuroplasticity": Sup. Fig.

### Supplementary Materials

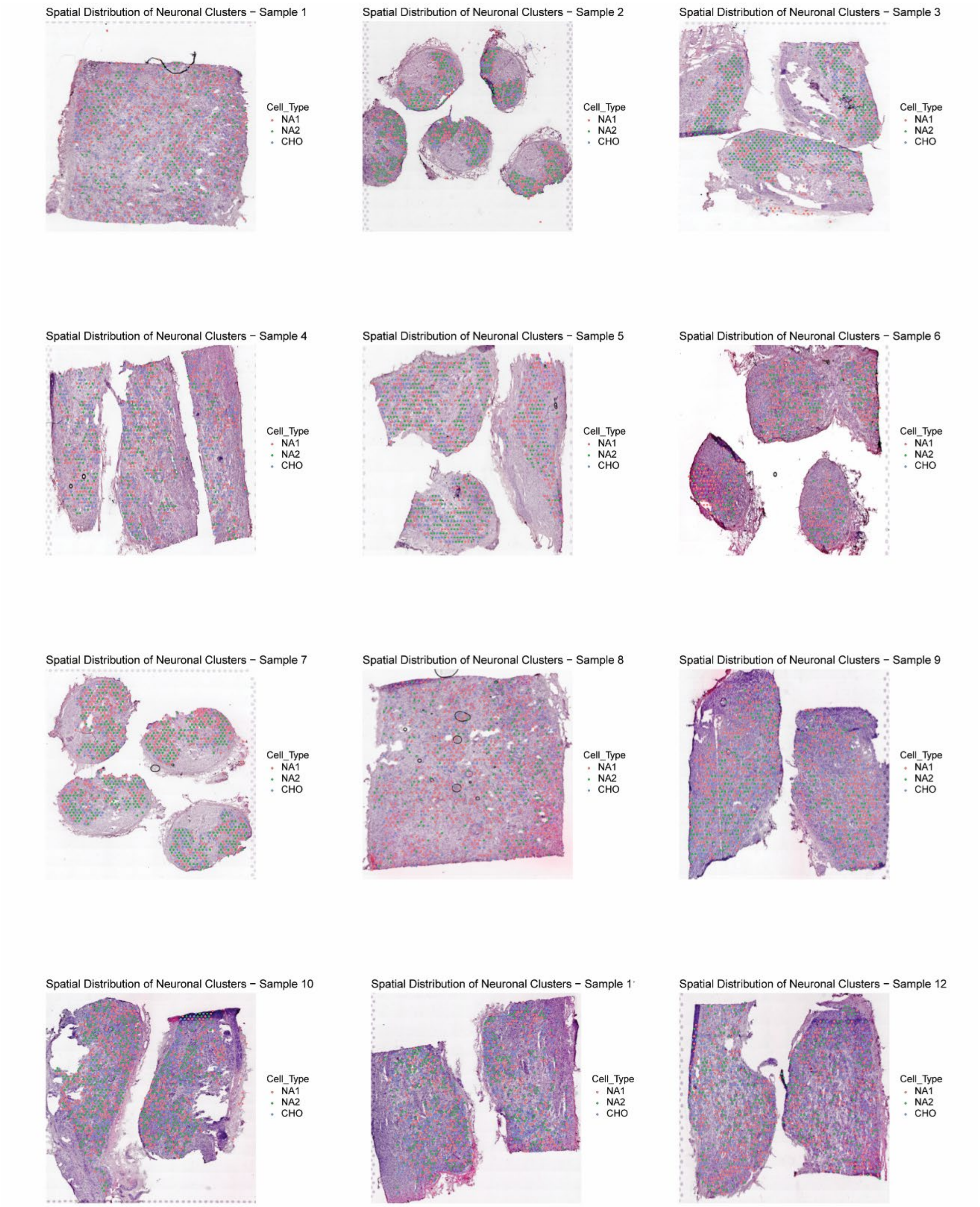


**Supplementary Figure S1. Neuronal spatial clusters across samples 1-12.** Visualization of neuronal spatial clusters within tissue sections. Overlapping colored barcodes indicate specific neuronal cell types localized within the core region of the sympathetic chain ganglia (SCG).

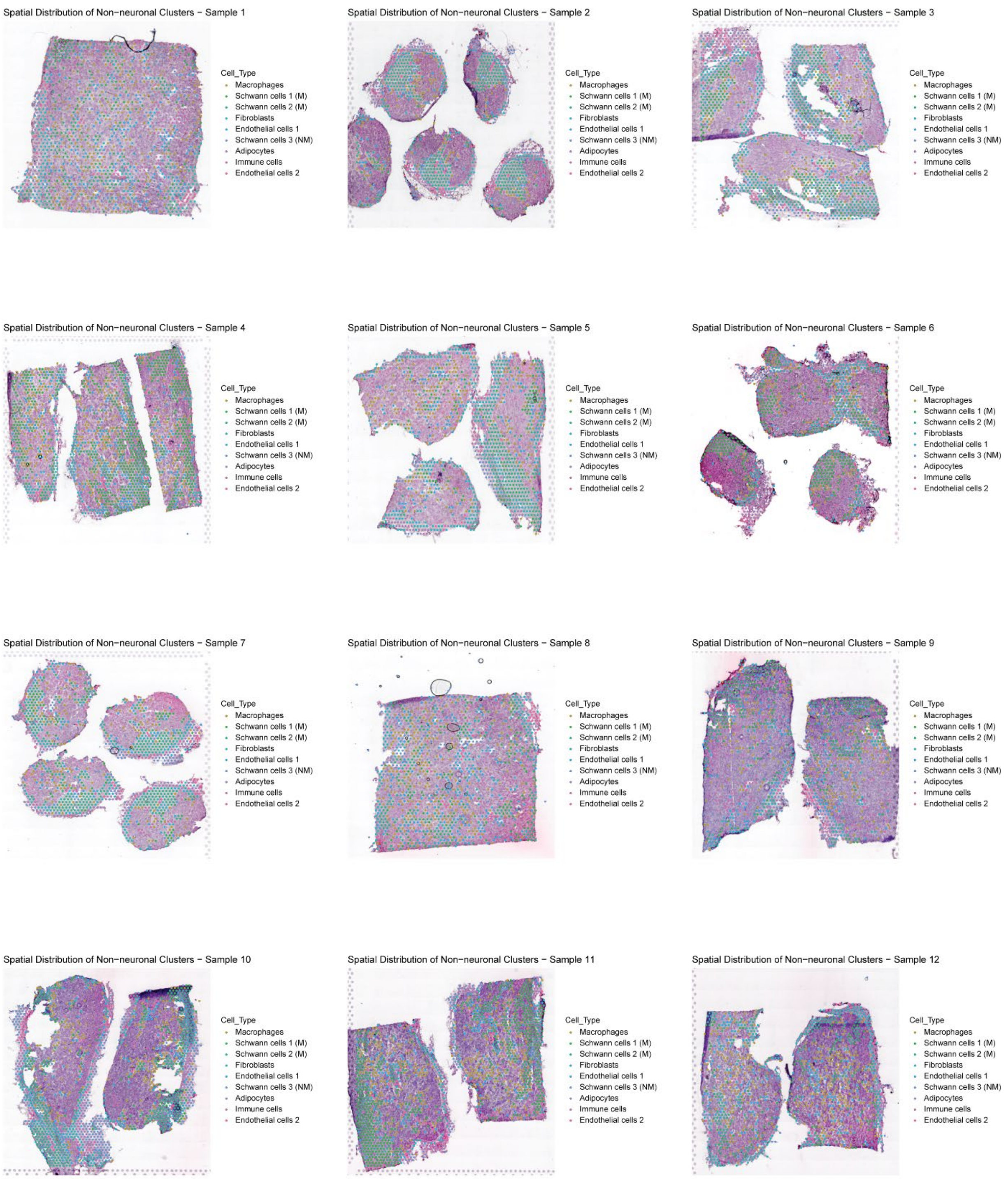


**Supplementary Figure S2. Non-neuronal spatial clusters across samples 1-12.** Visualization of non-neuronal spatial clusters within tissue sections. Overlapping colored barcodes indicate specific non-neuronal cell types localized surrounding the neuronal core of the sympathetic chain ganglia (SCG).

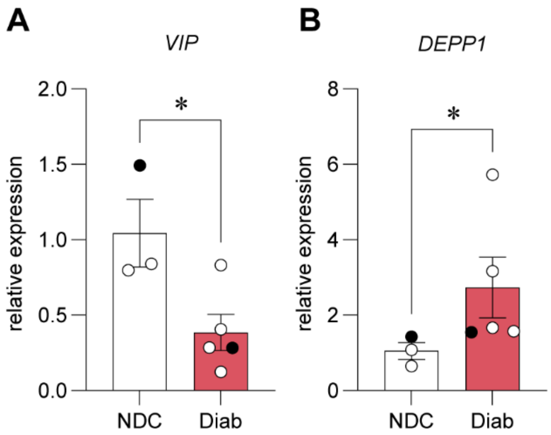


**Supplementary Figure S3. qPCR validation of spatial transcriptomics.** Using a separate cohort of SCGs obtained from different donors, we validated the change in expression of (A) *VIP* and (B) *DEPP1* mRNA. Black dots represent SCGs from samples used in the spatial transcriptomic experiment. White dots represent new samples. (A) *p<0.05, two-tailed t-test, (B) *p<0.05, Mann-Whitney test.

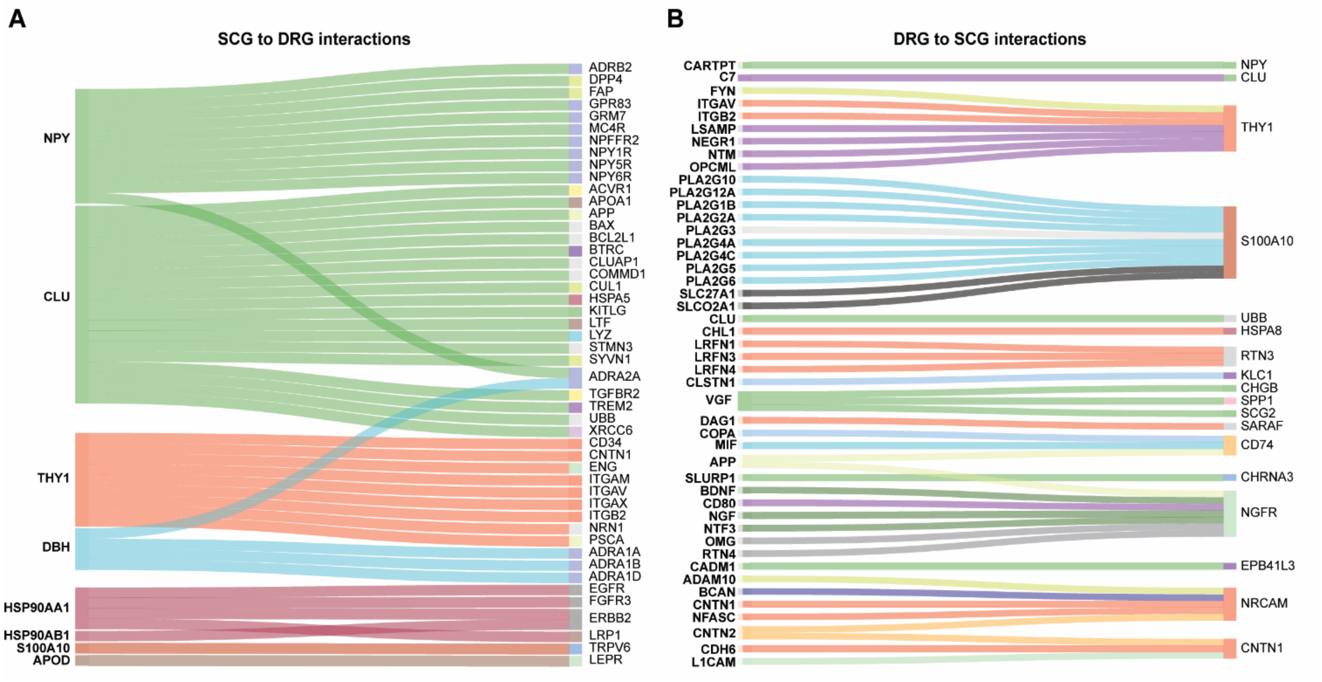


**Supplementary Figure S4. Molecular interaction analysis of Sympathetic Chain Ganglia (SCG) and dorsal root ganglion (DRG). A.** 8 SCG genes interact with 48 DRG genes, and **B.** 46 DRG genes interact with 18 SCG genes, with interactions represented by same-color curved lines. Complete input gene list in **Sup. Table 15.**

#### Supplementary methods

*Immunohistochemistry & Imaging*

We followed a modified immunocytochemistry protocol based on Optimum Cutting Temperature (OCT) embedded fresh frozen SCGs were cryosectioned at 20 μm (-20°C), and fixed with 10% buffered formalin (Thermo Fisher Scientific, 23245684) for 10 minutes. Sections underwent graded ethanol dehydrations (50%, 70%, 100% and 100%) before blocking for 1 hour. Primary antibodies were applied overnight at 4°C (anti-TH,1:200, ThermoFisher, MA1-24654; anti-Peripherin, 1:1000, EnCor Biotechnology, CPCA-Peri; anti-DBH, 1:200 ThermoFisher, MA5-46943). Secondary antibodies at 1:2000 (goat anti-rabbit Alexa Fluor 555, Thermo Fisher Scientific; goat anti-chicken Alexa Fluor 488, Thermo Fisher Scientific, A-11039; goat anti-mouse, Invitrogen, A21235) were incubated for 1 hour at room temperature, followed by DAPI nuclei staining (1:1000, Cayman Chemical Company, 14285).
Imaging was performed using an FV3000 confocal microscope (Evident Scientific) at 20x magnification, with brightness and contrast adjusted in Olympus CellSens software (v1.18).

*Human Sympathetic Chain Ganglia permeabilization optimization for 10X genomics Visium spatial transcriptomics*

Following the 10X Genomics Tissue Optimization User Guide (CG000238), 10μm-thick tissue sections were mounted onto the fiducial frames of Visium optimization slides and incubated for 12, 18, 24, 30, 36, and 42 minutes to optimize permeabilization time. Fluorescent signal intensity from neurons indicated RNA release efficiency, with the strongest signal at 30 minutes, identifying it as the optimal duration.

*Sequencing statistics*

Our sequencing run yielded a total of 1,027,706,540 reads, with an average of 96.21% successfully mapped to the reference human genome (GRCh38) and 98.85% identified as valid Unique Molecular Identifiers (UMIs). On average, we detected 20,949 genes across 2,209 spots under tissue. The mean number of reads per barcode was 36,049, while the median UMI counts per spot reached 2,820. More details in **Sup. Table 2.**

*Cell proportion analysis per Condition*

We extracted the number of cells per condition and population in R. The percentages were then calculated and visualized using Graphpad PRISM. **Sup. Tables 5 and 6.**

*Gene overlap identification in neurons (Venn Diagram)*

We identified differentially expressed genes (DEGs) for the NA1, NA2 and CHO populations using the `FindAllMarkers` function in Seurat with a log fold-change threshold of 0.25. Genes with $p < 0.05$ were filtered in Excel, and gene sets were compiled per cell type. The VennDiagram function in R was used to determine the number of overlapping genes between populations. **Sup. Tables 3 and 4**.

*Gene presence-absence comparisons in SCG and DRG*

To generate Table 1, we extracted the list of genes and their respective raw counts from a previously published human neuronal DRG dataset  and from our SCG neuronal populations. Genes were categorized as either completely absent or present (**Sup Tables 12-14**). Raw counts per gene within each tissue were converted to Counts Per Million (CPM) using the formula:

$$CPM = (SCG\ raw\ count\ /\ DRG\ raw\ count) \times 10^6$$

Lists were filtered to exclude mitochondrial genes, ribosomal proteins, and long non-coding RNAs. For genes expressed in both tissues, we calculated the fold change and identified the top five differentially expressed genes per tissue type. Since no statistical methods were employed for these comparisons, batch effects may be present.

$$Fold\ Change\ =\ (CPM\ SCG\ /CPM\ DRG)$$

*Interactome analysis between SCG and DRG*

We extracted the average expression data for all genes from the previously built neuronal susbset using the AverageExpression function, stratified by cell type. A CSV file was generated containing gene expression data filtered for $p < 0.05$ in the NA1, NA2, and CHO populations. This file was then uploaded to SensoryOmics to generate a Sankey plot (**Sup. Table 15**) (1). The analysis was performed twice using the human DRG Visium dataset available on the platform—once with DRG as the ligand and once as the receptor.

*RNA Extraction and RT-qPCR*

SCGs were flash-frozen on dry ice and embedded in OCT, then sectioned at 150 µm using a Leica CM1950 cryostat. Excess OCT was manually trimmed using a sterile razor blade. Tissue sections were transferred to 2 mL Precellys lysing tubes (Bertin Corp, Cat# P000918-LYSK0-A) pre-chilled on dry ice. Homogenization was performed in the same tubes using 900 µL of QIAzol Lysis Reagent (Qiagen, Cat# 79306). Disruption was executed using the Minilys homogenizer (Bertin Technologies) at the lowest available setting, with a 30-second burst protocol in a 4°C cold room. RNA isolation was performed on ice with the RNeasy Plus Universal Mini Kit (Qiagen, Cat# 73404) in accordance with the manufacturer's protocol, within a designated RNase-free sterile environment. RNA concentrations were quantified using the RNA-specific quantification settings of a NanoDrop 2000 spectrophotometer (ThermoScientific). For reverse transcription (RT), cDNA synthesis was carried out using the iScript Advanced cDNA Synthesis Kit for RT-qPCR (Bio-Rad, Cat# 1725038), adhering to the manufacturer's standardized protocol, with reactions executed on a Bio-Rad T100 thermal cycler. Quantification of synthesized cDNA was performed using the single-stranded DNA (ssDNA) settings of the NanoDrop 2000 spectrophotometer (ThermoScientific). Quantitative PCR (qPCR) was conducted with the SsoAdvanced Universal SYBR Green Supermix (Bio-Rad, Cat# 1725271) on Applied Biosystems 7500 Real-Time PCR System, following the manufacturer's instructions. $RT^2$ qPCR primer assays (Qiagen, 330001) were obtained to quantify transcript levels of human VIP (RefSeq: NM_003381, GeneGlobe ID: PPH02072A) and human DEPP1 (RefSeq: NM_007021, GeneGlobe ID: PPH10525A) normalized to human GAPDH (RefSeq: NM_002046, GeneGlobe ID: PPH00150F) and human HPRT1(RefSeq: NM_000194, GeneGlobe ID: PPH01018C). Data analysis was performed using the comparative CT method in Microsoft Excel and GraphPad Prism.
